# Enhancing microalgal productivity through bioactive substances, light, and CO_2_

**DOI:** 10.1101/2025.11.26.690871

**Authors:** Andrea Llanes, Wiston Quiñones, Natalia Herrera

## Abstract

Microalgae play a crucial role in ecosystems, from oxygen production to sustaining food webs and offering valuable applications in industry and environmental management. Increasing their productivity in terms of biomass yield, cultivation time, and nutritional or metabolite quality remains a major challenge. This study assessed the effects of 10 bioactive substances (including lactones, phytohormones, and natural extracts), four light wavelengths, and four CO_2_ injection regimes on *Arthrospira platensis*, *Chlorella vulgaris*, *Ankistrodesmus falcatus*, and *Tetradesmus dimorphus*, using linear modeling and LSD (Least Significant Difference) post hoc tests. N-butyryl-DL-homoserine lactone significantly enhanced *A. platensis* growth (94.4%), while naphthaleneacetic acid and indole-3-butyric acid promoted *T. dimorphus* growth (138.1% and 115.5%, respectively). More accessible alternatives, such as *Aloe vera* and coconut water, also stimulated growth: *A. platensis*, *C. vulgaris*, and *A. falcatus* increased by 85.3%, 69.2%, and 87.7% with *Aloe vera*, while *T. dimorphus* increased 80.5% with coconut water. Regarding light quality, red light (600–700 nm) benefited *A. platensis* and *A. falcatus* (49.2% and 20.8%, respectively), whereas blue light (400–490 nm) favored *C. vulgaris* and *T. dimorphus* (57.7% and 31.5%, respectively). CO_2_ injection further enhanced biomass production and carbon fixation, particularly in *C. vulgaris* and *A. falcatus* (73.5% and 53.5%, respectively). However, combined treatments did not produce additive effects, suggesting complex interactions. Overall, these findings demonstrate the potential of bioactive substances and environmental conditions to improve microalgal performance and highlight the importance of investigating synergistic effects and scalability for large-scale production.

## 1. Introduction

Microalgae can synthesize value-added products such as polysaccharides, proteins, lipids, and biologically active compounds that have important applications in the pharmaceutical, cosmetic, food, and biofuel production industries [1]. In addition, they play a crucial role in maintaining ecosystems, mitigating metal toxicity, preserving aquatic and terrestrial biodiversity, treating wastewater, and, in a general sense, capturing CO_2_ [2]. Microalgae biomass is a renewable and efficient resource that can be key to meeting the growing demand for sustainable alternatives, especially in sectors such as bioenergy, food security, and reducing the environmental footprint [3]. Due to this significant potential to play a significant role in various areas and industries, there has been considerable interest in optimizing microalgal biomass production.

Optimizing microalgal biomass production is crucial for its scalable and economically viable use [4]. One way to do this is by controlling some of the growth factors such as pH, temperature, salinity, nutrients, light intensity and quality, and the addition of inducing substances, among others [5,6]. Chunzhuk et al. [7] reported that increasing light intensity along with the addition of CO_2_ can improve the production of *Arthrospira platensis, Chlorella ellipsoidea, Chlorella vulgaris, Gloeotila pulchra,* and *Elliptochloris subsphaerica*. Meanwhile, Xie et al. [8] y Seemashree et al. [9] reported that the addition of inducing substances such as phytohormones in the culture medium can also improve the production of *Chlorella vulgaris* (cv-31), *Porphyridium purpureum*, and *Dunaliella salina*.

Despite these advances, most studies have evaluated growth factors individually, with limited attention to their combined effects or to the use of alternative, low-cost inducers. Understanding how different factors interact is crucial, as responses are often species-specific and non-linear, and outcomes cannot be predicted from single-factor experiments alone. In particular, exploring the potential of natural extracts as sustainable bioactive substances, along with variations in light quality and CO_2_ enrichment, could provide valuable insights for both fundamental physiology and applied large-scale cultivation [10,11]. Such approaches may help identify strategies that are not only effective in enhancing biomass productivity but also economically and environmentally feasible for industrial implementation [12].

Given the variability in species-specific responses, it is necessary to investigate how individual strains react to different factors and their interactions. Accordingly, this study aimed to assess the effects of 10 bioactive substances, four light wavelengths, and four CO_2_ injection times—individually and in combination—on the growth and/or nutritional quality of *Chlorella vulgaris*, *Ankistrodesmus falcatus*, *Tetradesmus dimorphus*, and the cyanobacterium *Arthrospira platensis*, with the goal of improving biomass production.

## 2. Materials and methods

### 2.1. Obtaining and identifying microalgae

The cyanobacterium *Arthrospira platensis* was acquired from the Algae Culture Laboratory - Biology Department of the National University of Colombia. *Ankistrodesmus falcatus* was obtained from the Laboratory for Environmental Health Assessment and Promotion, Oswaldo Cruz Institute (Fiocruz), Brazil; *Chlorella vulgaris* and *Tetradesmus dimorphus* were isolated from different soils around the water reservoir in the village of La Palma in Carmen de Viboral, Antioquia, Colombia. To isolate the microalgae, 10 g of each soil sample containing *Chlorella* and *Tetradesmus* were first weighed and transferred to 400 mL of BBM culture medium. The samples were kept on a shelf at room temperature for 3 days for conditioning.

Then, 50 mL of each culture was taken and centrifuged (Sigma 2-16P Universal Centrifuge, Sigma Laborzentrifugen GmbH, Osterode am Harz, Germany) at 5000 rpm (251 x g) for 5 minutes to concentrate the microalgae and remove unwanted particles. The precipitate obtained was resuspended in 400 mL of culture medium specific to each microalga and left to grow for 7 days. To clean the cultures of possible contaminants, three transfers were performed every 7 days, in which 10 mL of the previous culture was taken and finally transferred to 400 mL of new culture medium.

### 2.2. Microalgae cultivation and maintenance

Once cleaned and fully adapted, the cultures were maintained as follows: 500 mL Erlenmeyer flasks were used, each containing 400 mL of culture medium. The flasks were fitted with a rubber stopper containing two ports: one for connecting the aeration tube and another to release internal pressure. Aeration was continuous and maintained at 1.0 L/min. The culture media used were Zarrouk medium for *A. platensis*, BBM medium for *C. vulgaris* and *T. dimorphus*, and ASM-1 medium for *A. falcatus*.

The culture conditions were as follows: a 12 h:12 h light/dark photoperiod, temperature of 25 (±1) °C, light intensity of 100 µmol m⁻² s⁻¹, constant aeration at 1.0 L/min, pH 7 (±0.5) for BBM and ASM-1 media, and pH 9 (±0.5) for Zarrouk medium. Additionally, the culture media were renewed every 15 days.

For taxonomic identification, the morphological characteristics of each microalga were observed under a microscope (Nikon Eclipse E-200, Nikon Corporation, Tokyo, Japan), considering their structural features (size, organization, shape, pigments, motility) [13–15] following the Algaebase [16].

### 2.3. Exposure of microalgae to ten bioactive substances

Each microalga was exposed for 12 days to ten different bioactive substances, as detailed in **Table 1** and illustrated in **Fig 1**. The assays were conducted under the culture conditions described in section 2.2.

**Fig 1.**
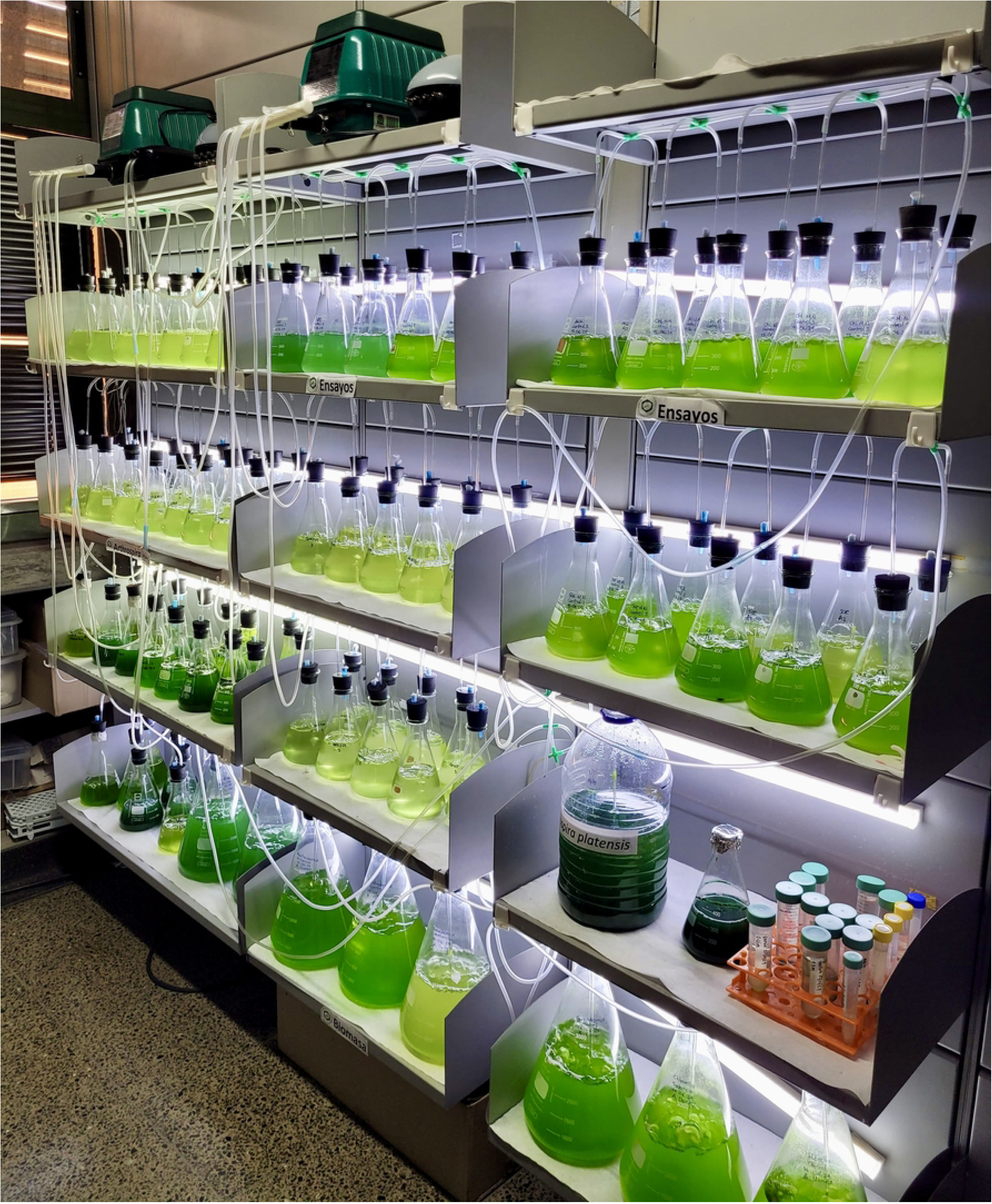
Experimental setup of the bioactive substance exposure assays.

**Table 1.**
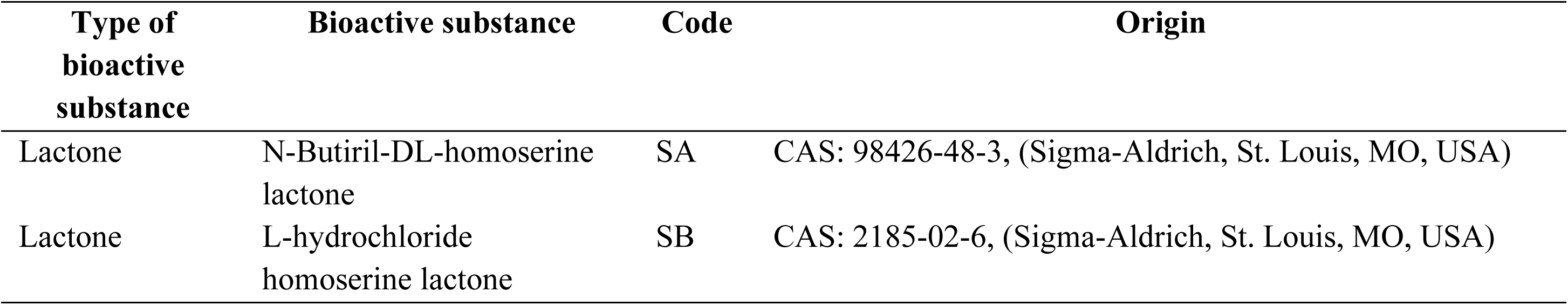
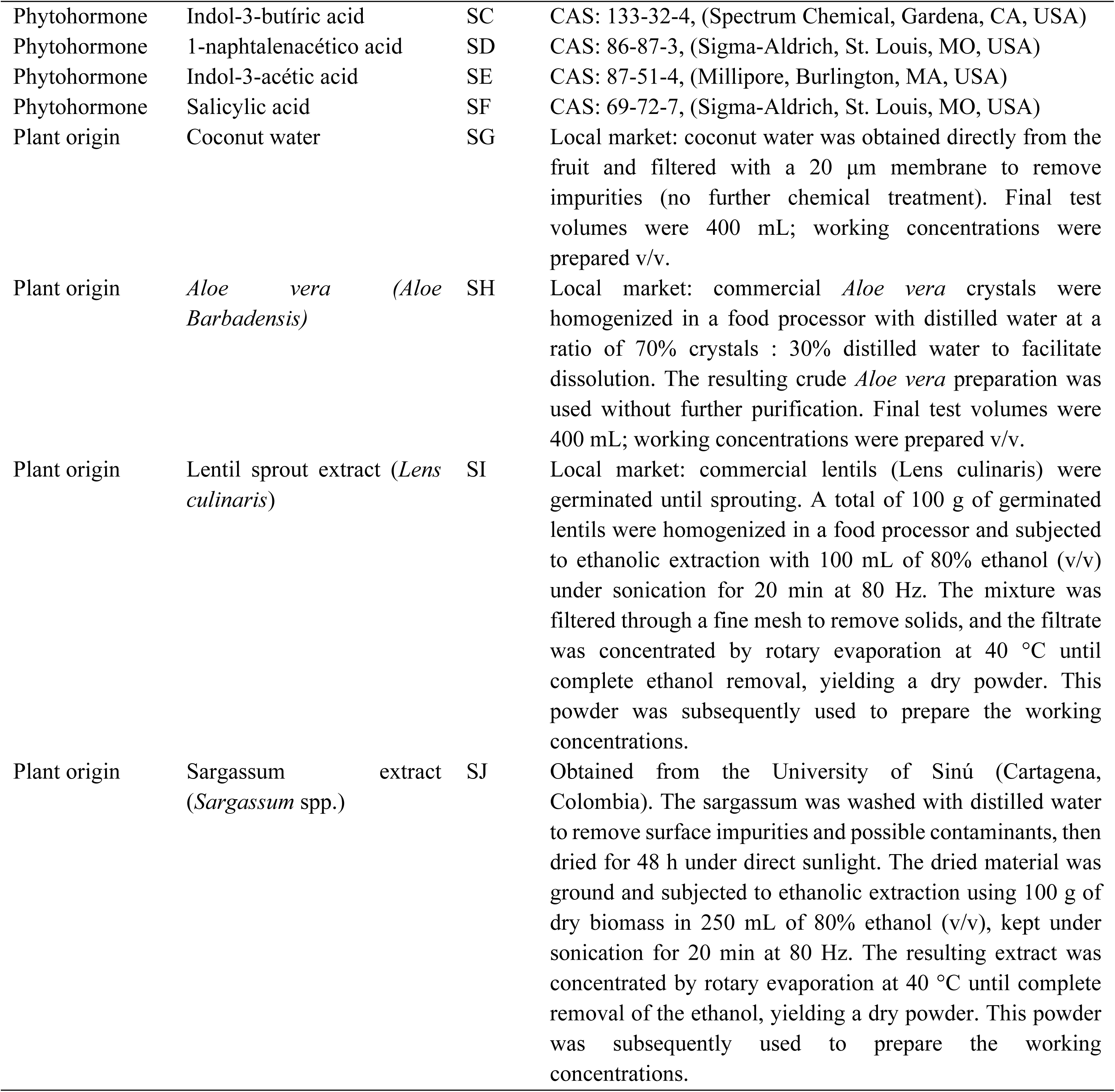
Bioactive substances used as potential inducers of microalgae growth evaluated.

| Type of bioactive substance | Bioactive substance | Code | Origin |
| --- | --- | --- | --- |
| Lactone | N-Butiril-DL-homoserine lactone | SA | CAS: 98426-48-3, (Sigma-Aldrich, St. Louis, MO, USA) |
| Lactone | L-hydrochloride homoserine lactone | SB | CAS: 2185-02-6, (Sigma-Aldrich, St. Louis, MO, USA) |
| Phytohormone | Indol-3-butíric acid | SC | CAS: 133-32-4, (Spectrum Chemical, Gardena, CA, USA) |
| Phytohormone | 1-naphtalenacético acid | SD | CAS: 86-87-3, (Sigma-Aldrich, St. Louis, MO, USA) |
| Phytohormone | Indol-3-acético acid | SE | CAS: 87-51-4, (Millipore, Burlington, MA, USA) |
| Phytohormone | Salicylic acid | SF | CAS: 69-72-7, (Sigma-Aldrich, St. Louis, MO, USA) |
| Plant origin | Coconut water | SG | Local market: coconut water was obtained directly from the fruit and filtered with a 20 µm membrane to remove impurities (no further chemical treatment). Final test volumes were 400 mL; working concentrations were prepared v/v. |
| Plant origin | *Aloe vera* (*Aloe Barbadensis*) | SH | Local market: commercial *Aloe vera* crystals were homogenized in a food processor with distilled water at a ratio of 70% crystals : 30% distilled water to facilitate dissolution. The resulting crude *Aloe vera* preparation was used without further purification. Final test volumes were 400 mL; working concentrations were prepared v/v. |
| Plant origin | Lentil sprout extract (*Lens culinaris*) | SI | Local market: commercial lentils (*Lens culinaris*) were germinated until sprouting. A total of 100 g of germinated lentils were homogenized in a food processor and subjected to ethanolic extraction with 100 mL of 80% ethanol (v/v) under sonication for 20 min at 80 Hz. The mixture was filtered through a fine mesh to remove solids, and the filtrate was concentrated by rotary evaporation at 40 °C until complete ethanol removal, yielding a dry powder. This powder was subsequently used to prepare the working concentrations. |
| Plant origin | Sargassum extract (*Sargassum* spp.) | SJ | Obtained from the University of Sinú (Cartagena, Colombia). The sargassum was washed with distilled water to remove surface impurities and possible contaminants, then dried for 48 h under direct sunlight. The dried material was ground and subjected to ethanolic extraction using 100 g of dry biomass in 250 mL of 80% ethanol (v/v), kept under sonication for 20 min at 80 Hz. The resulting extract was concentrated by rotary evaporation at 40 °C until complete removal of the ethanol, yielding a dry powder. This powder was subsequently used to prepare the working concentrations. |

Lactones, phytohormones, lentil sprout extract, and sargassum extract were used at final concentrations: C1 = 0.1 μg/mL, C2 = 1.0 μg/mL, C3 = 5.0 μg/mL, and C4 = 10.0 μg/mL. Four concentrations were also tested for coconut water and *Aloe vera*: C1 = 1%, C2 = 3%, C3 = 7%, and C4 = 10% v/v. All treatments were applied in a final culture volume of 400 mL (in 500 mL Erlenmeyer flasks). A negative control containing only culture medium was included. Each assay was performed in triplicate.

### 2.4. Exposure to different wavelengths

Each microalga was exposed to four different wavelengths for 12 days. The assays were conducted under the culture conditions described in section 2.2, except with a continuous photoperiod of 24:0 h (light/dark) to maximize growth rate and biomass productivity, and to isolate the effect of light spectrum as the experimental variable while avoiding the influence of light–dark cycles [17–19]. To achieve this, the inner walls of the cultivation chambers were lined with reflective mirror film, and a commercial RGB LED strip with Bluetooth control was used to adjust each target wavelength. Based on manufacturer specifications (≈300–500 lm m⁻¹, 4–6 m total strip length per chamber) and enclosure geometry, the photon flux density was estimated to be in the range of 80–150 µmol m⁻² s⁻¹. The wavelengths tested were: blue light (400–490 nm, L1), red light (600–700 nm, L2), green light (490–550 nm, L3), and yellow light (570–580 nm, L4). Additionally, a white light control (400–700 nm) was included. All experiments were conducted in triplicate.

### 2.5. Exposure of microalgae to four CO_2_ injection times

Each microalga was exposed to CO_2_ at a flow rate of 1.0 L/min (99.95%, Messer SE & Co. KGaA) at times T1=30 s, T2=60 s, T3=90 s, and T4=120 s, for 12 days, every 24 hours, under the culture conditions described in section 2.2. Additionally, a control without the addition of CO_2_ was used; each test was performed in triplicate. The final biomass of each culture was harvested by sedimentation for *C. vulgaris, T. dimorphus,* and *A. falcatus*, and using a 15 μm filter mesh for *A. platensis*. The biomass was washed with Milli-Q water, freeze-dried (LABCONCO FreeZone 12L, Labconco Corporation, Kansas City, MO, USA), and the dry weight was determined on a balance.

### 2.6. Evaluation of microalgae cell growth

The effect of each of the 10 bioactive substances, wavelengths, and CO_2_ on microalgae growth was measured using four calibration curves made for each microalgae species. The curves were constructed as follows: Starting with a known number of cells at time zero, the progress of the culture was monitored with a Multiskan Spectrum (Thermo Scientific, Waltham, MA, USA) at 630 nm. During the first 15 days, measurements were taken every 24 hours, and from day 16 to day 30, they were taken every 48 hours. In addition, counts were performed in a Neubauer chamber (Marienfeld, Lauda-Königshofen, Germany) for *C. vulgaris*, *T. dimorphus*, and *A. falcatus*, and in a Sedgewick Rafter counting chamber (Pyser Optics, Edenbridge, UK) for *A. platensis*. Then, a calibration curve was constructed from which the equation relating the variables absorbance and cells/mL was obtained.

To evaluate the effect of the 10 bioactive substances, wavelengths, and CO_2_ on microalgae growth, three samples were taken from each culture: one on day zero, as initial data, and two at the end of each experimental test, after 12 days, as final data. Of the two final samples, one was used to quantify cell growth by spectrophotometry, applying the equation obtained from each calibration curve. For this purpose, 300 μL of each culture was transferred to a 96-well plate, and the absorbance was measured using a wavelength reader [20]. Each measurement was performed in triplicate. The second sample was used for microscoPGI observations with a microscope (Nikon, Eclipse E-200, Tokyo, Japan) to evaluate possible changes in cell morphology (S1 File. Cell morphology).

In addition, the percentage of growth induction (PGI) of each microalga exposed to each bioactive substance, wavelength, and CO_2_ was calculated using the following equation [20]:

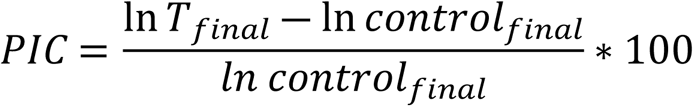

Where T = Treatment, corresponds to the densities of cells exposed and not exposed (control) to the different bioactive substances, and ln = Natural logarithm.

Finally, CO_2_ fixation (Fi) was calculated from the dry weight of the microalgae biomass and the carbon content of each species (% C), determined in the total oxidizable organic carbon analyses, using the following equation (adapted from Wu et al. [21]):

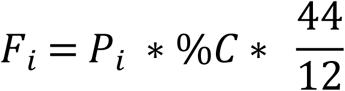

Where Fi is the CO_2_ fixation for day i and Pi is the productivity obtained for day i, in discontinuous cultures.

### 2.7. Evaluation of the effect of various factors (bioactive substances, wavelength, and CO_2_) and their combinations

Based on the results obtained in sections 2.3, 2.4 and 2.5 (bioactive substances, light wavelengths, and CO_2_, respectively), the effects of these factors—considered inducers of growth and biomass production in microalgae according to their individual outcomes—were evaluated both individually and in combination. To this end, the four microalgal species were exposed to seven culture conditions for 12 days under the general conditions described in section 2.2. Growth was measured spectrophotometrically, as previously described (section 2.6), using in each case the optimal parameters determined in the preceding experiments. The seven conditions evaluated were:

- Control: Standard culture conditions (section 2.2)
- Condition 1 (C1): Bioactive substance
- Condition 2 (C2): Light wavelength
- Condition 3 (C3): CO_2_ exposure time
- Condition 4 (C4): Bioactive substance, light wavelength, and CO_2_ exposure time
- Condition 5 (C5): Bioactive substance and light wavelength
- Condition 6 (C6): Bioactive substance and CO_2_ exposure time
- Condition 7 (C7): Light wavelength and CO_2_ exposure time

### 2.8. Analysis of the nutritional composition of the biomass of each microalga

For the optimal treatments identified in the results of section 2.7, scaling was performed in 3 L Erlenmeyer flasks of 2.5 L, using 400 mL of culture from a 12-day stock of each microalga. The control consisted of scaling with constant aeration at 1 L/min, maintaining the culture conditions described in section 2.2. After 30 days of culture, the biomass produced was collected. The final biomass of each culture was harvested by sedimentation for *C. vulgaris, T. dimorphus,* and *A. falcatus*, and using a 15 μm filter mesh for *A. platensis*. The biomass was washed with Milli-Q water, freeze-dried (LABCONCO FreeZone 12L, Labconco Corporation, Kansas City, MO, USA) (pre-frozen at −20 °C for 24 h; operating vacuum 0.2–0.4 mbar), and the dry weight was determined on a balance.

The lyophilized biomass of each cultivated microalga was analyzed for carbon, nitrogen, and mineral content. Total oxidizable organic carbon was determined using the Oxidant-Titrimetric method (NTC 5167); nitrogen was analyzed by the volumetric Kjeldahl method (ISO 5983); phosphorus was quantified using the UV–VIS spectrophotometric method (NTC 4981); and minerals including calcium, copper, iron, magnesium, manganese, potassium, sodium, and zinc were analyzed using the atomic absorption spectrophotometric method (NTC 5151). It is worth noting that these mineral elements are not synthesized by microalgae but originate from the culture medium; however, their accumulation in the biomass may reflect species-specific differences in uptake and bioaccumulation capacity, which is of interest from both nutritional and biotechnological perspectives [22].

### 2.9. Statistical analysis

Initially, the data were normalized using the natural logarithm to use the same scale for all data. A linear model was created, and an ANOVA (Analysis of Variance) was applied with a significance level of 0.05. The assumptions of normality (Kolmogorov-Smirnov test), independence (Durbin Watson test), and homoscedasticity (Breusch Pagan test) were validated. After performing ANOVA tests, significant differences were found, so it was decided to use the post hoc LSD (Least Significant Difference) test for multiple comparison analysis. All tests were analyzed at a significant level of 0.05 and a confidence level of 95%. For all analyses, RStudio 9.2 statistical software was used, with R 4.3 processing, Core Team (2023). The data were analyzed using a linear model, which showed a good fit in all cases according to the R² values and met most of the assumptions (Table A in S3 File).

## 3. Results

### 3.1. Effect of the evaluated factors (bioactive substances, wavelength, and CO_2_) and their combinations on the growth of *Arthrospira platensis*

Initially, it was found that 1.0 μg/mL of N-Butyryl-DL-homoserine lactone (SA) has a positive effect on the growth of *A. platensis*, with a PGI=94.4% and statistically significant differences compared to the other bioactive substances (p=3.87e-12, Fig 1A). This statistical significance (letter “a” according to the LSD post hoc test) is shared with other treatments, such as *Aloe vera* (SH) at 3% (p=6.38e-12) with a PGI=91.9% and 7% *Aloe vera* (SH) (p=3.23e-11) with a PGI=88.1%. In addition, exposure to red light (600-700 nm, L2) generated an appreciable increase in growth, with a PGI of 49.2%, with statistically significant differences compared to the other wavelengths (p=0.000125; Fig 1B). Finally, CO_2_ injection (60 s, T2) also promoted significant growth, reaching a PGI of 41.2% (p=1.12e-07; Fig 1C). In this case, 0.230 g of CO_2_ was fixed, representing an increase of 0.068 g compared to the control (0.162 g).

Based on the results, *Aloe vera* was chosen for further testing as an inducing substance because it is a more accessible alternative compared to N-Butyryl-DL-homoserine lactone. Although the latter showed the highest effect (PGI 94.4%), *Aloe vera* at 3% achieved a PGI of 91.9%, with no statistically significant differences between the two. In evaluating the effect of the factors and their combinations, 3% *Aloe vera* (Condition 1) was statistically significant compared to the other conditions (p=<2.14e-10, Fig 2). Furthermore, Table 2 shows that the nutritional profile of *A. platensis* presented slight variations under the effect of 3% *Aloe vera*, with increases in calcium, phosphorus, iron and zinc content.

**Fig 2.**
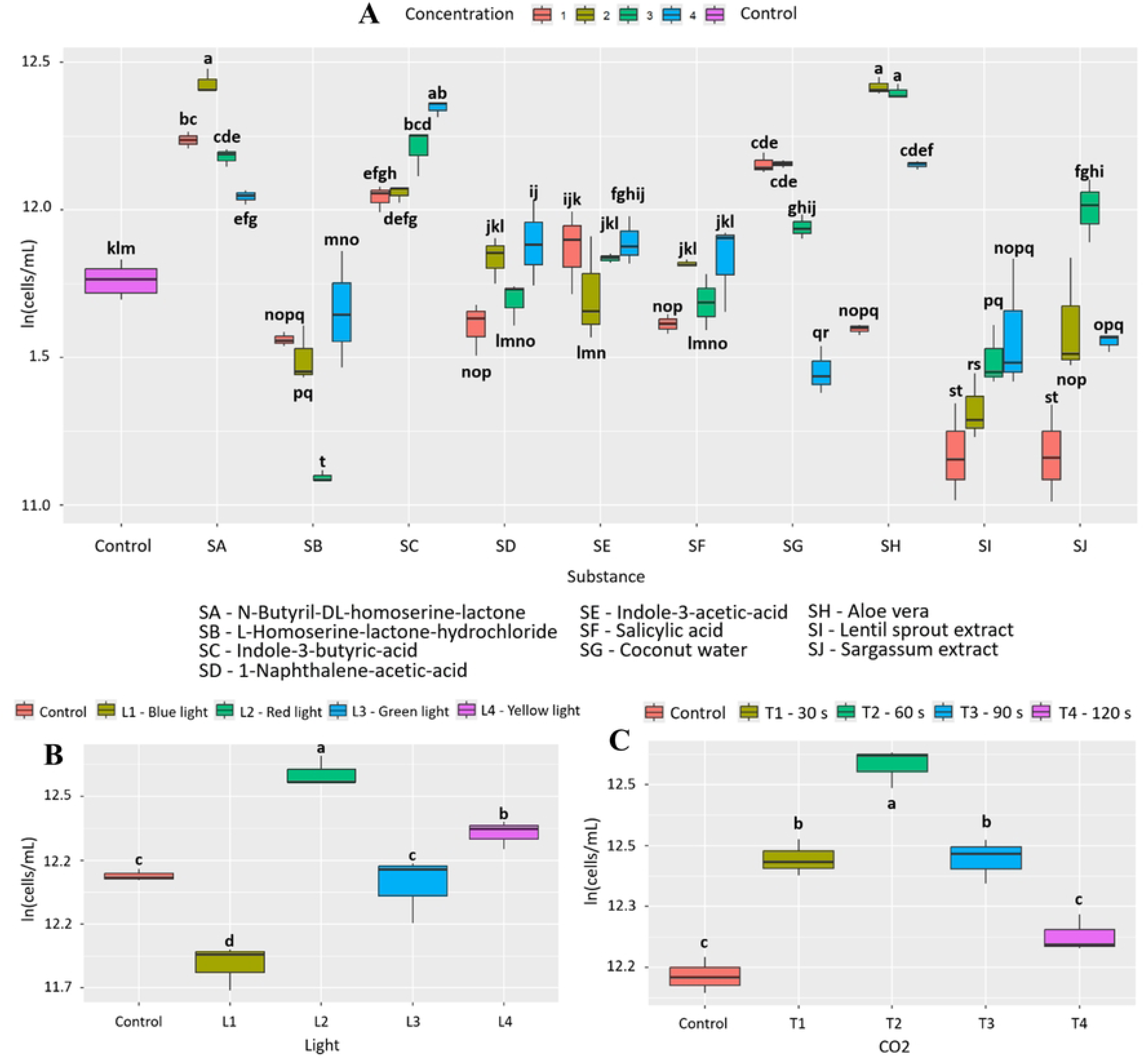
Effect of the factors evaluated on the growth of *A. platensis*. (A) 10 bioactive substances evaluated at 4 concentrations (B) 4 wavelengths and (C) 4 CO_2_ injection times. Letters indicate statistically significant differences between treatments (p < 0.05).

**Table 2.** Carbon, nitrogen, and mineral content of the best culture conditions for each microalga.

| Analysis | *A. platensis* |  | *C. vulgaris* |  | *A. falcatus* |  | *T. dimorphus* |  |
| --- | --- | --- | --- | --- | --- | --- | --- | --- |
|  | Control | *Aloe vera* 3% | Control | *Aloe vera* 3% | Control | *Aloe vera* 1% | Control | Coconut water 3% |
| Total oxidizable organic carbon | 34.3 % | 34.3% | 34.3% | 38% | 45.0% | 46.3% | 34.0% | 33.1% |
| Calcium (mg/kg) | 19 | 190 | 190 | 181 | 40 | 39 | 197 | 345 |
| Cooper (mg/kg) | ND | <5 | <5 | <5 | ND | ND | <5 | <5 |
| Phosphorous (mg/kg) | <100 | 142 | 142 | 123 | <100 | <100 | 171 | 458 |
| Iron (mg/kg) | <5 | 8 | 8 | 5 | <5 | <5 | 8 | 9 |
| Magnesium (mg/kg) | <50 | <50 | <50 | <50 | <50 | <50 | 50 | 101 |
| Manganese (mg/kg) | <5 | <5 | <5 | <5 | <5 | <5 | <5 | <5 |
| Nitrogen (g/100g) | 10.9 | 9.2 | 6.9 | 6.1 | 5.7 | 5.9 | 5.5 | 5.3 |
| Potassium (g/100g) | 0.67 | 0.56 | 0.56 | 0.60 | 0.55 | 0.64 | 0.58 | 0.61 |
| Sodium (mg/kg) | <500 | <500 | <500 | <500 | <500 | <500 | <500 | <500 |
| Zinc (mg/kg) | ND | 15 | 15 | 11 | ND | ND | 15 | 21 |

ND: not detected

### 3.2. Effect of materials and conditions (bioactive substances, wavelength, and CO_2_) and their combinations on the growth of *Chlorella vulgaris*

Initially, it was found that 3% *Aloe vera* (SH) has a positive effect on the growth of *C. vulgaris*, with a PGI=76.2% and statistically significant differences compared to the other bioactive substances (p<2.0e-16, Fig 3A). This statistical significance is shared with other treatments such as L homoserine lactone (SB) at 10.0μg/mL (p=<2.0e-16) with a PGI=76.0% and coconut water (SG) at 1% (p=<2.0e-16) with a PGI=73.6%. In addition, exposure to blue light (400-490 nm, L1) generated a notable increase in growth, with a PGI=57.7% with statistically significant differences compared to the other wavelengths (p = 2.47e-08; Fig 3B). Finally, CO_2_ injection (60s, T2) also promoted significant growth, reaching a PGI of 73.5% (p=1.67e-08; Fig 3C). In this case, 0.384 g of CO_2_ was fixed, representing an increase of 0.166 g compared to the control (0.218 g).

**Fig 3.**
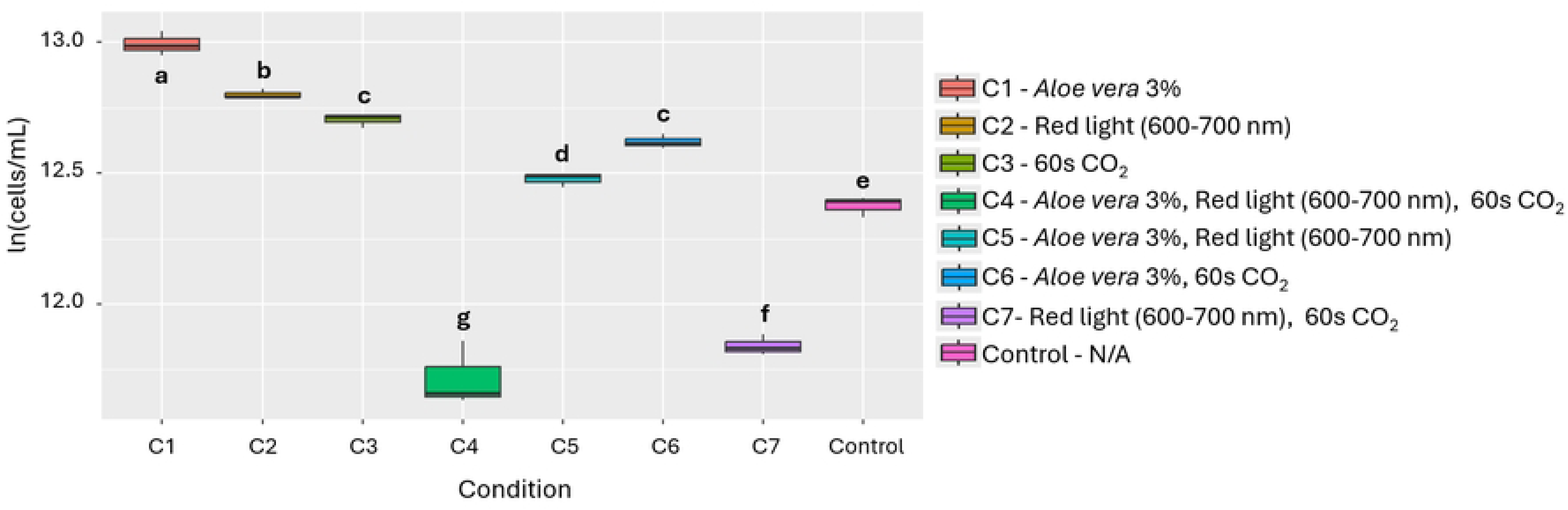
Effect of the individual and combined use of the 3 evaluated factors on the growth of *A. platensis*. Different letters indicate statistically significant differences among treatments (p < 0.05).

In evaluating the effect of individual materials and conditions, as well as their combinations, 3% *Aloe vera* (Condition 1) was statistically significant compared to the other conditions (p=2.46e-12, Fig 4). In addition, Table 2 shows that the nutritional profile of *C. vulgaris* exhibited a increase in total oxidizable organic carbon and potassium with 3% *Aloe vera*, accompanied by slight decreases in calcium, phosphorus, iron, and zinc content.

**Fig 4.**
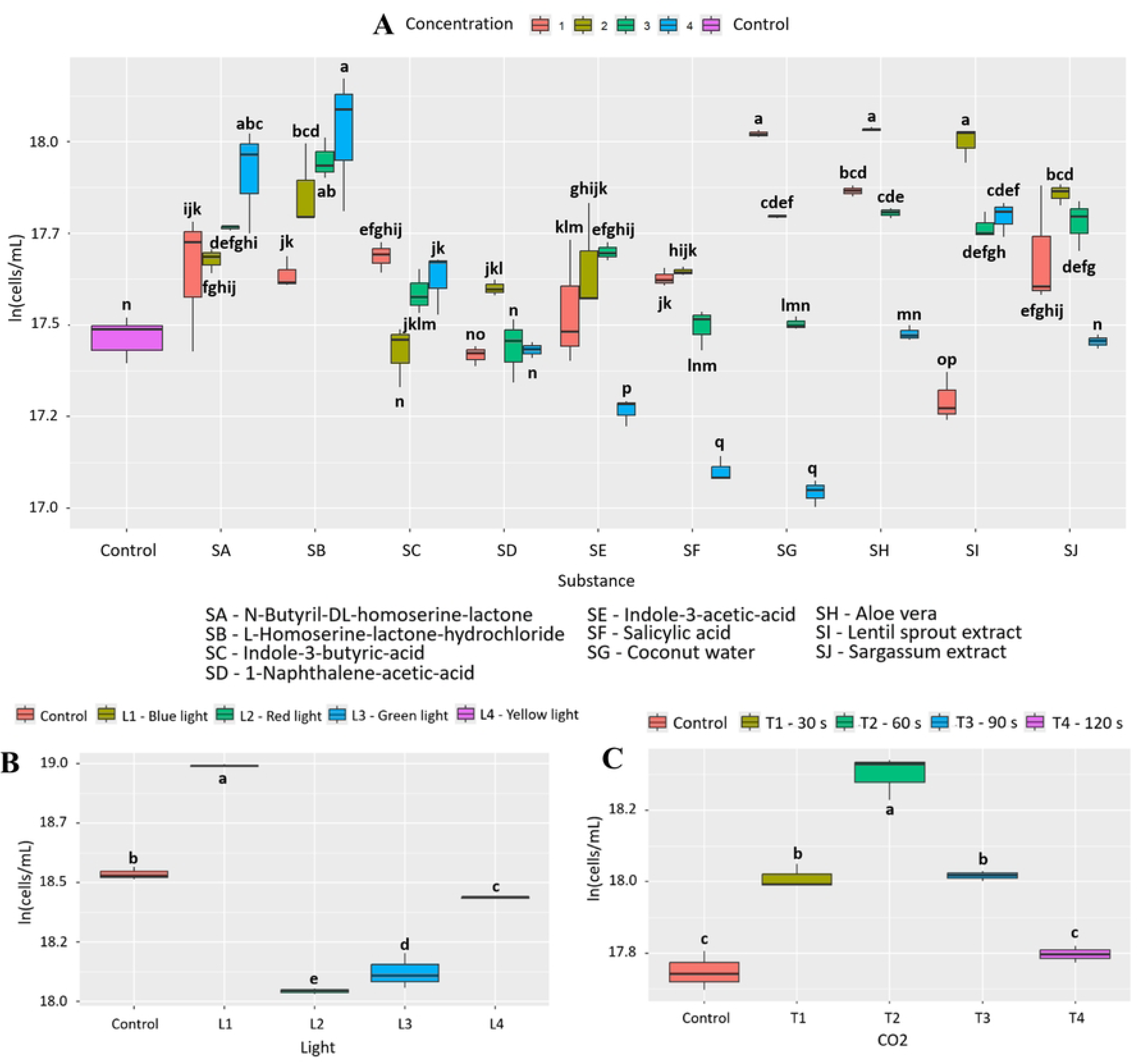
Effect of the factors evaluated on the growth of *C. vulgaris*. (A) 10 bioactive substances evaluated at 4 concentrations (B) 4 wavelengths and (C) 4 CO_2_ injection times. Letters indicate statistically significant differences between treatments (p < 0.05).

### 3.3. Effect of materials and conditions (bioactive substances, wavelength, and CO_2_) and their combinations on the growth of *Ankistrodesmus falcatus*

Initially, it was found that 1% *Aloe vera* (SH) has a positive effect on the growth of *A. falcatus*, with a PGI=97.4% and statistically significant differences compared to the other bioactive substances (p=2.0e-16, Fig 5A). This statistical significance is shared with other treatments such as 1% coconut water (SG) (p=2.22e-13) with a PGI=76.6% and 3% *Aloe vera* (SH) (p=5.93e-13) with a PGI=74.3%. In addition, exposure to red light (600-700 nm, L2) generated a slight increase in growth, with a PGI of 20.8%, with statistically significant differences compared to the other wavelengths (p=0.00177; Fig 5B). Finally, CO_2_ injection (30s, T1) also promoted high growth, reaching a PGI of 53.5% (p=6.38e-08, Fig 5C). In this case, 0.271 g of CO_2_ was fixed, representing an increase of 0.101 g compared to the control (0.154 g).

**Fig 5.**
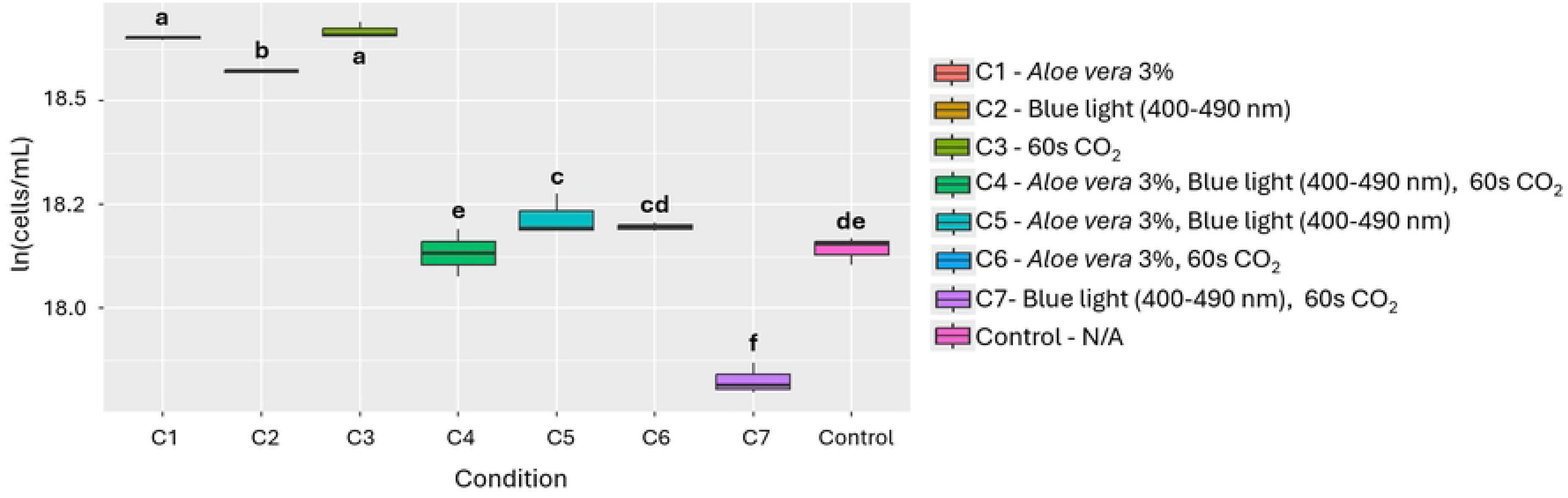
Effect of the individual and combined use of the 3 evaluated factors on the growth of *C. vulgaris*. Different letters indicate statistically significant differences among treatments (p < 0.05).

In evaluating the effect of materials and conditions, as well as their combinations, 1% *Aloe vera* (Condition 1) was statistically significant compared to the other conditions (p=3.95e-15, Fig 6). Furthermore, Table 2 shows that the nutritional profile of *A. falcatus* displayed minor increases in total oxidizable organic carbon and potassium content under the effect of 1% *Aloe vera*.

**Fig 6.**
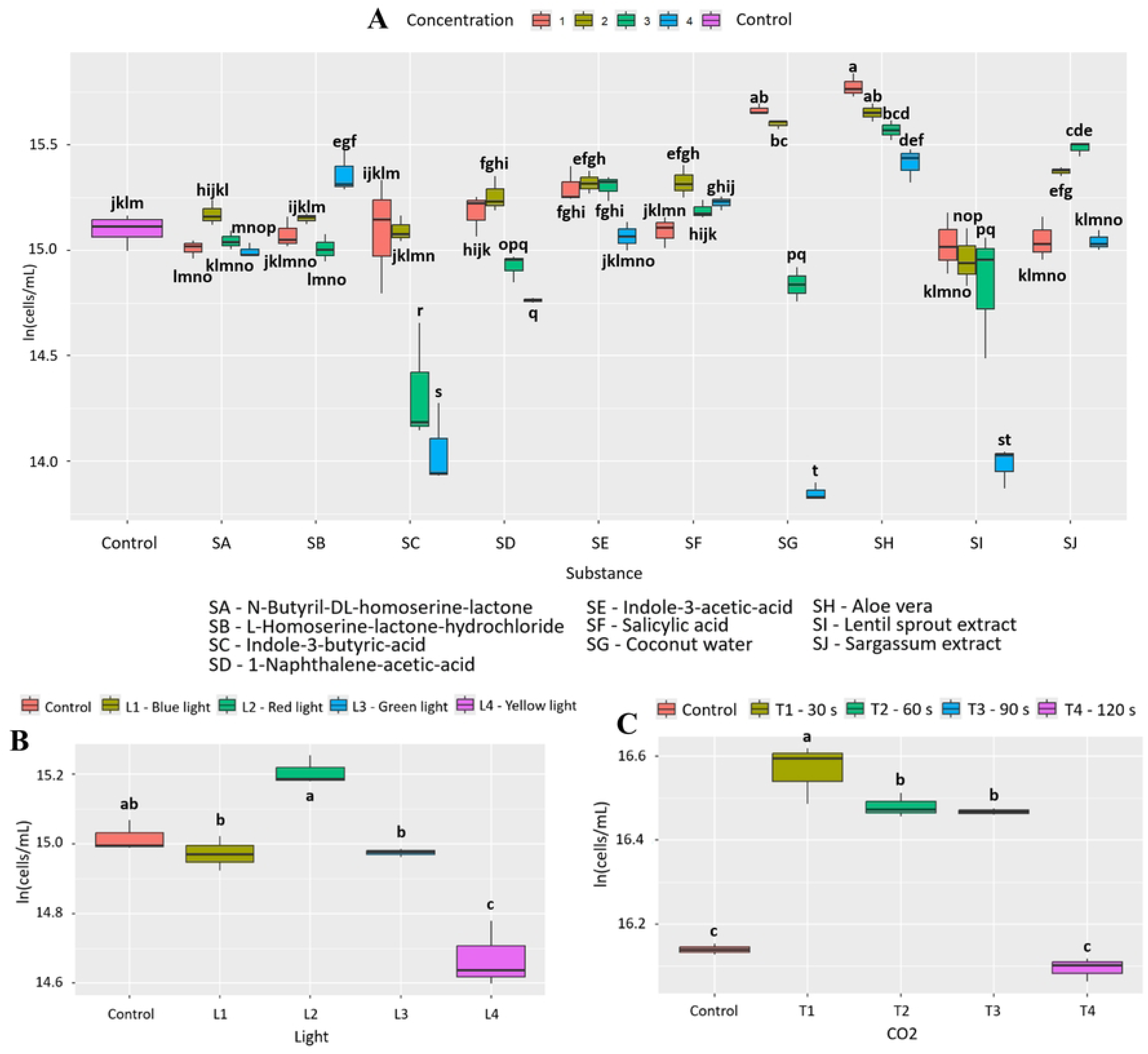
Effect of the factors evaluated on the growth of *A. falcatus*. (A) 10 bioactive substances evaluated at 4 concentrations (B) 4 wavelengths and (C) 4 CO_2_ injection times. Letters indicate statistically significant differences between treatments (p < 0.05).

### 3.4. Effect of materials and conditions (bioactive substances, wavelength, and CO_2_) and their combinations on the growth of *Tetradesmus dimorphus*

Initially, it was found that 10.0 μg/mL of 1-naphthaleneacetic acid (SD) has a positive effect on the growth of *T. dimorphus*, with a PGI=138.1% and statistically significant differences compared to the other bioactive substances (p=2.39e-16, Fig 7A). This statistical significance is shared with other treatments such as indole-3-butyric acid (SC) 10.0 μg/mL (p=1.91e-13) with a PGI=115.5% and coconut water (SG) 3% (p=1.90e-11) with a PGI=93.8%. Additionally, exposure to blue light (400-490 nm, L2) generated an appreciable increase in growth, with a PGI of 31.5%, with statistically significant differences compared to the other wavelengths (p=1.01e-06; Fig 7B). Finally, CO_2_ injection (60 s, T2) slightly promoted growth, reaching a PGI of 21.7% (p=0.000824, Fig 7C). In this case, 0.196 g of CO_2_ was fixed, representing an increase of 0.043 g compared to the control (0.096 g).

**Fig 7.**
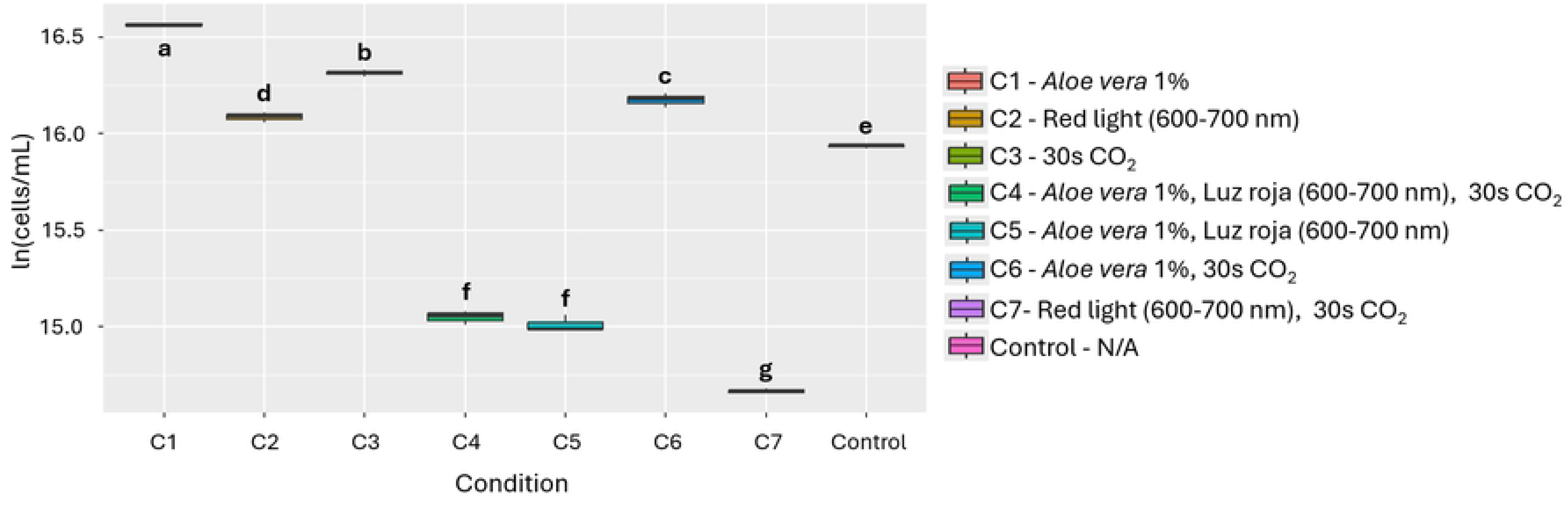
Effect of the individual and combined use of the 3 evaluated factors on the growth of *A. falcatus*. Different letters indicate statistically significant differences among treatments (p < 0.05).

Based on the results and like aloe, coconut water (SG) (PGI=93.8%) was chosen as an inducer because it is a more accessible alternative to other bioactive substances such as 1-naphthaleneacetic acid (SD, 138.1%) and indole-3-butyric acid (SC, 115.5%). Although its effect on growth is lower, it still shows a significant difference (p=1.90e-11), which justifies its selection in this study. In the evaluation of the effect of the factors and their combinations, 3% coconut water (Condition 1) was statistically significant compared to the other conditions (p=2.50e-10, Fig 8). In addition, Table 2 shows that the nutritional profile of *T. dimorphus* presented increases in calcium, phosphorus, iron, magnesium, potassium, and zinc content, accompanied by a slight decrease in oxidizable organic carbon content.

**Fig 8.**
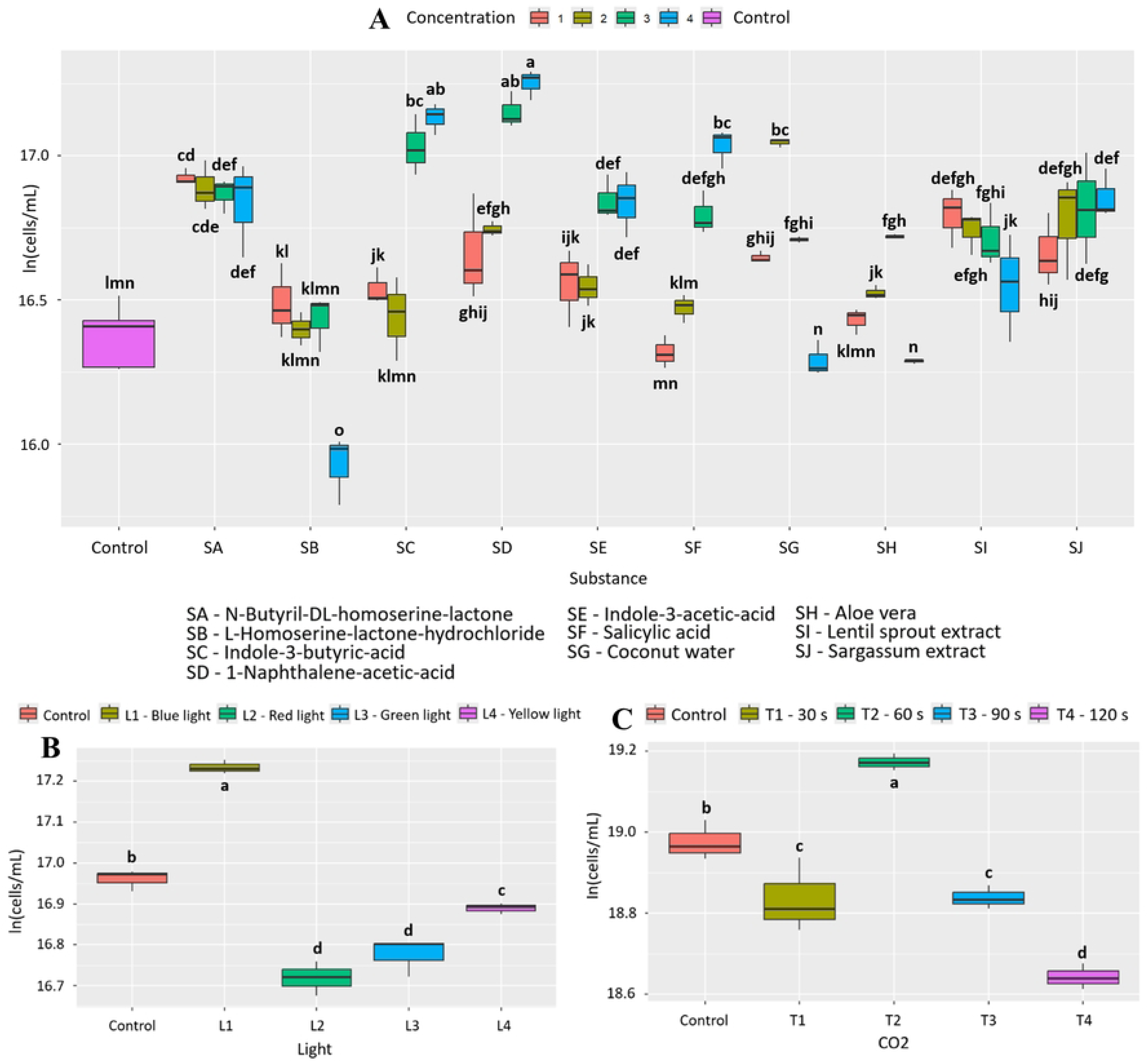
Effect of the factors evaluated on the growth of *T. dimorphus*. (A) 10 bioactive substances evaluated at 4 concentrations (B) 4 wavelengths and (C) 4 CO_2_ injection times. Letters indicate statistically significant differences between treatments (p < 0.05).

**Fig 9.**
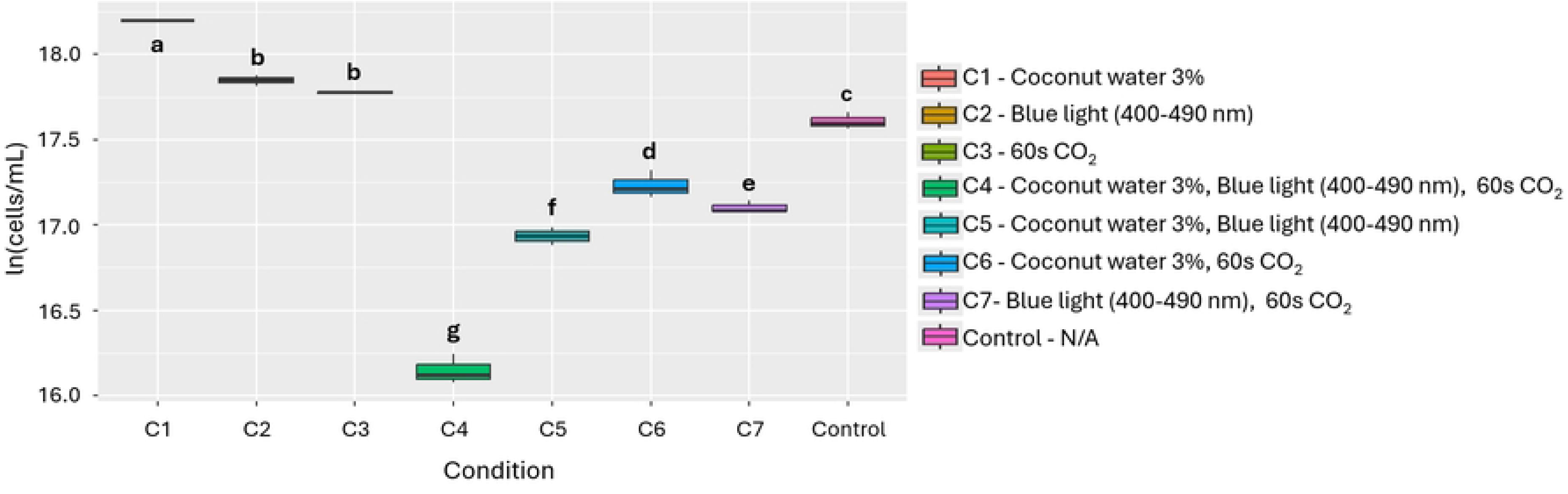
Effect of the individual and combined use of the 3 evaluated factors on the growth of *T. dimorphus*. Different letters indicate statistically significant differences among treatments (p < 0.05).

## 4. Discussion

This study evaluated the impact of different cultivation conditions on *Arthrospira platensis, Chlorella vulgaris, Ankistrodesmus falcatus,* and *Tetradesmus dimorphus* to optimize their growth and/or nutritional quality. Conditions such as the use of bioactive substances, wavelength, and CO_2_ injection were analyzed, allowing the optimal conditions for each species to be identified.

The conditions evaluated in this study were selected based on their potential to induce microalgae growth and improve biomass quality. In particular, phytohormones and lactones were used, compounds widely recognized for their ability to regulate key physiological processes in plants and microalgae. Studies have shown that phytohormones such as indole-3-acetic acid, salicylic acid, indole-3-butyric acid, and naphthalene acetic acid, among others, can stimulate cell growth and the production of metabolites such as proteins, lipids, chlorophylls, and antioxidants [23–25]. Likewise, lactones (especially N-acylhomoserine lactones) have been studied as growth inducers in various microalgae species, probably due to their role in intercellular communication and the modulation of physiological responses [20,26,27]; which were shown to have an inducing effect on the growth of different microalgae studied.

In addition, natural extracts such as *Aloe vera* and coconut water were included due to their previous use, attributed to their content of cytokinins, vitamins, sugars, and other bioactive compounds [28,29]. On the other hand, the spectral quality of light directly influences photosynthesis and microalgae growth. In previous studies, red and blue light are particularly effective in increasing biomass in various species [30]. Finally, CO_2_ injection was considered as a strategy to increase the availability of inorganic carbon, which can improve the photosynthetic rate and promote faster growth [31].

One of the most relevant findings of this study is the effectiveness of widely available, low-cost, natural substances, such as *Aloe vera* and coconut water, in significantly improving the growth of the microalgae tested. Regarding nutritional composition (oxidizable carbon and nitrogen content), the results showed minimal variations between treatments, with no significant changes compared to the control. Both are raw materials that are accessible in various regions of the world, especially in tropical contexts, which facilitates their integration into cultivation systems without requiring expensive technologies or inputs that are difficult to obtain. This accessibility, coupled with their biodegradable nature and traditional use in food and agricultural applications reported in the literature, positions these biostimulants as sustainable alternatives to synthetic or more specialized compounds [32,33]. Together, these results not only reinforce the development of industrial solutions with low environmental impact and economic viability, especially in sectors such as aquaculture, agriculture, and water treatment. Together, these results not only reinforce the development of industrial solutions with low environmental impact and economic viability, especially in sectors such as aquaculture, agriculture, and water treatment.

The addition of *Aloe vera* proved to be an effective strategy for stimulating growth, with particularly notable effects on *A. platensis* (PGI=88.9%), *C. vulgaris* (PGI=75.0%), and *A. falcatus* (PGI=85.2%) (Figs 2, 4, 6 and Figs A, C and C in S2 File). This effect can be explained by the presence of bioactive compounds in the extract, such as polysaccharides, vitamins, antioxidants, and minerals, which have been associated with promoting cell proliferation and improving photosynthetic efficiency [34–36]. In addition to growth, changes in nutritional composition were observed (Table 2), such as increases in calcium, phosphorus, iron and zinc in *A. platensis*, total oxidizable organic carbon and potassium in *A. falcatus* and *C. vulgaris*. However, decreases in certain micronutrients, such as calcium, phosphorus, iron, and zinc in *C. vulgaris*, were also recorded, which could indicate internal nutrient redistribution processes or possible competition in their absorption under specific cultivation conditions [22].

Coconut water, on the other hand, showed a notable effect on the growth of *T. dimorphus*, especially at 3%, with a PGI of 80.5% (Fig 8 and Fig D in S2 File), surpassing *Aloe vera* in this species (PGI=39.8%). This behavior could be due to its rich composition of cytokinins and essential nutrients such as potassium, calcium, and phosphorus, which stimulate key physiological processes such as cell division and photosynthesis [37]. Nutritional improvements were also observed, with increases in calcium, phosphorus, iron, magnesium, potassium, and zinc. However, in other species such as *C. vulgaris* and *A. falcatus*, *Aloe vera* was more efficient, highlighting the need to select biostimulants based on the target species. Other bioactive substances, such as N-Butyryl-DL-homoserine lactone in *A. platensis*, L-Homoserine-lactone hydrochloride in *C. vulgaris*, and plant growth regulators such as naphthaleneacetic acid and indole-3-butyric acid in *T. dimorphus*, showed high efficacy as microalgae growth inducers (Figs 1A, 3A, 7A and Table B in S3 File).

Although the main effect observed was on growth stimulation, the nutritional changes detected were relatively modest and not always consistent across species. This outcome is not unexpected, as enhanced biomass production does not necessarily imply major shifts in cellular composition. In many cases, rapid cell proliferation primarily increases the total amount of biomass available, while the nutrient content per cell remains stable [38,39]. From a biotechnological perspective, this result is still highly relevant, since higher biomass yields are often the primary objective in large-scale microalgae cultivation, and even slight improvements in mineral composition can add value to the final product. Therefore, the findings of this study confirm that while biostimulants such as *Aloe vera* and coconut water can enhance growth efficiency, their nutritional effects may be more subtle and species-dependent.

On the other hand, spectral lighting had a significant and differentiating effect on microalgae growth, highlighting the need to adjust light according to the pigment physiology of each species. Red light (600–700 nm) was most effective in *A. platensis* and *A. falcatus*, with increases of 49.2% and 20.8%, respectively (Figs 1B, 5B, and Table C in S3 File), possibly due to their content of phycobiliproteins such as phycocyanin, which absorb efficiently in that range [40,41]. In contrast, *C. vulgaris* and *T. dimorphus* responded better to blue light (400–490 nm), with increases of 57.7% and 31.5% (Figs 3B, 7B and Table C in S3 File), attributable to their higher proportion of chlorophylls a and b, optimized to capture this spectrum [42,43]. This adaptation could be reinforced by mechanisms such as the xanthophyll cycle or lower sensitivity to photoinhibition, which favors their productivity under light intensity [44].

CO_2_ supply proved to be an effective stimulus for growth and carbon fixation in the microalgae evaluated, although with different responses depending on the species. *A. platensis, C. vulgaris,* and *T. dimorphus* responded favorably to a 60-second injection (Figs 1C, 3C, 7C and Table D in S3 File), suggesting that intermediate exposures optimize photosynthesis without generating harmful accumulations of dissolved CO_2_ [45]. On the other hand, *A. falcatus* showed its highest yield with only 30 seconds (Fig 5C and Table D in S3 File), indicating remarkable efficiency in short-term carbon uptake. These differences indicate the need to adjust CO_2_ injection time according to the species, prioritizing not only biomass yield, but also the efficiency in the assimilation of inorganic carbon [46].

Finally, the combination of stimuli such as bioactive substances, light spectra, and CO_2_ injection times does not always produce additive or synergistic effects on microalgae growth (Figs 2, 4, 6, 8 and Table E in S3 File). In fact, in species such as *A. platensis* and *A. falcatus*, some combinations (such as condition C6, which includes *Aloe vera* and CO_2_) showed moderate increases in growth, but were outperformed by individual treatments. Conversely, in species such as *C. vulgaris* and *T. dimorphus*, the combination of materials and conditions often resulted in significant reductions in growth. All this suggests that the simultaneity of stimuli may generate physiological interference or stress effects that inhibit the positive response observed in isolation.

All this indicates that each microalga responds differently and that the interaction between them must be carefully evaluated before scaling up their use in production systems [47,48]. Rather than seeking multiple combinations, it might be more efficient to design specific treatments tailored to the physiological characteristics of each species [49]. Future research should focus on the combined optimization of these materials and conditions, exploring synergies, evaluating their scalability in large-scale production systems, and investigating the underlying biochemical mechanisms to develop more efficient and sustainable strategies for microalgae cultivation.

From a biotechnological perspective, *T. dimorphus* and *A. falcatus* emerge as the most promising species, not only due to their greater responsiveness to individual stimuli, but also due to their nutritional profiles rich in minerals and organic carbon. *T. dimorphus*, in particular, combines accelerated growth with a favorable nutritional profile, while *A. falcatus* stands out for its carbon fixation efficiency under optimal conditions. These characteristics make them attractive candidates for food, nutraceutical, and environmental mitigation applications.

## 5. Conclusions

Analysis of the four microalgae species evaluated, *Arthrospira platensis, Chlorella vulgaris, Ankistrodesmus falcatus,* and *Tetradesmus dimorphus*, showed significant improvements, primarily in growth under optimal conditions, with no significant changes in nutritional composition compared to the control. The implementation of biostimulants, particularly natural extracts such as *Aloe vera* and coconut water, specific wavelengths, and CO_2_ injection strategies, proved effective in enhancing microalgae growth, although the response was not always positive when several factors were applied simultaneously. These results highlight the value of integrating biotechnological tools with a detailed understanding of species-dependent responses, where increased biomass is emerging as an initial step for future research aimed at industrial and environmental applications.

## Acknowledgments

This work was supported by Minciencias (Colombia) (COD 1115- 914-91810, Ct 111-2022). We extend our gratitude to Professor Fernando Echeverri for his support during the research and data analysis.

## **7.** Supporting information

S1 File. Cell morphology

S2 File. 30 day growth curves

S3 File. Supplementary data tables

## References

1. Rodríguez-Roque MJ, Flores-Córdova MA, Salas-Salazar NA, Soto-Caballero MC, Valdivia-Nájar CG, Sánchez-Vega R. Microalgae as source of bioaccessible and bioavailable compounds. Handbook of Food and Feed from Microalgae [Internet]. Elsevier; 2023 [cited 2023 Dec 19]. p. 519–27. 10.1016/B978-0-323-99196-4.00016-4

2. Sharma A, Arya SK. Ecological and environmental services of microalgae. Valorization of Microalgal Biomass and Wastewater Treatment [Internet]. Elsevier; 2023 [cited 2023 Dec 19]. p. 261–315. 10.1016/B978-0-323-91869-5.00007-7

3. Ahmad A, W. Hassan S, Banat F. An overview of microalgae biomass as a sustainable aquaculture feed ingredient: food security and circular economy. Bioengineered [Internet]. 2022 [cited 2023 Dec 19];13:9521–47. 10.1080/21655979.2022.2061148

4. Subramanian S, Sayre RT. The right stuff; realizing the potential for enhanced biomass production in microalgae. Front Energy Res [Internet]. 2022 [cited 2023 Dec 19];10:979747. 10.3389/fenrg.2022.979747

5. Pedruzi GOL, Amorim ML, Santos RR, Martins MA, Vaz MGMV. Biomass accumulation-influencing factors in microalgae farms. Rev bras eng agríc ambient [Internet]. 2020 [cited 2025 Jan 14];24:134–9. 10.1590/1807-1929/agriambi.v24n2p134-139

6. Khor JG, Lim HR, Chia WY, Chew KW. Automated Cultivation System for Microalgae: Growth Factors andControl. CNF [Internet]. 2022 [cited 2025 Jan 14];18:776–9. 10.2174/1573401318666220421132428

7. Chunzhuk EA, Grigorenko AV, Kiseleva SV, Chernova NI, Vlaskin MS, Ryndin KG, et al. Effects of Light Intensity on the Growth and Biochemical Composition in Various Microalgae Grown at High CO2 Concentrations. Plants [Internet]. 2023 [cited 2025 Jan 14];12:3876. 10.3390/plants12223876

8. Xie Z, Cao Y, Peng S, Zhang X, Kong W. Effects of six phytohormones on the growth behavior and cellular biochemical components of Chlorella vulgaris 31 [Internet]. 2023 [cited 2025 Jan 14]. 10.21203/rs.3.rs-2671883/v1

9. Seemashree MH, Chauhan VS, Sarada R. Phytohormone supplementation mediated enhanced biomass production, lipid accumulation, and modulation of fatty acid profile in Porphyridium purpureum and Dunaliella salina cultures. Biocatalysis and Agricultural Biotechnology [Internet]. 2022 [cited 2025 Jan 14];39:102253. 10.1016/j.bcab.2021.102253

10. López-Hernández JF, García-Alamilla P, Palma-Ramírez D, Álvarez-González CA, Paredes-Rojas JC, Márquez-Rocha FJ. Continuous Microalgal Cultivation for Antioxidants Production. Molecules [Internet]. 2020 [cited 2025 Aug 30];25:4171. 10.3390/molecules25184171

11. Wu K, Ying K, Liu L, Zhou J, Cai Z. High irradiance compensated with CO2 enhances the efficiency of Haematococcus lacustris growth. Biotechnology Reports [Internet]. 2020 [cited 2025 Aug 30];26:e00444. 10.1016/j.btre.2020.e00444

12. Tam LT, Ha NC, Thom LT, Zhu J, Wakisaka M, Hong DD. Ferulic acid extracted from rice bran as a growth promoter for the microalga Nannochloropsis oculata. J Appl Phycol [Internet]. 2021 [cited 2025 Aug 30];33:37–45. 10.1007/s10811-020-02166-5

13. Bellinger EG, Sigee DC. Freshwater algae: identification,enumeration and use as bioindicators. 2nd edition. Chichester, West Sussex ; Hoboken, NJ: John Wiley & sons Inc; 2015.

14. Gualtieri P, Barsanti L. Algae: Anatomy, Biochemistry, and Biotechnology [Internet]. 3rd ed. Boca Raton: CRC Press; 2022 [cited 2025 Jan 15]. 10.1201/9781003187707

15. John DM, Whitton BA, Brook AJ. The freshwater algal flora of the British isles: an identification guide to freshwater and terrestrial algae. Cambridge: Cambridge university press; 2002.

16. Guiry M, Guiry G. AlgaeBase [Internet]. National University of Ireland, Galway. 2024. https://www.algaebase.org/

17. Holdmann C, Schmid-Staiger U, Hornstein H, Hirth T. Keeping the light energy constant — Cultivation of Chlorella sorokiniana at different specific light availabilities and different photoperiods. Algal Research [Internet]. 2018 [cited 2025 Aug 20];29:61–70. 10.1016/j.algal.2017.11.005

18. Kato Y, Fujihara Y, Vavricka CJ, Chang J-S, Hasunuma T, Kondo A. Light/dark cycling causes delayed lipid accumulation and increased photoperiod-based biomass yield by altering metabolic flux in oleaginous Chlamydomonas sp. Biotechnol Biofuels [Internet]. 2019 [cited 2025 Aug 20];12:39. 10.1186/s13068-019-1380-4

19. Maroneze MM, Deprá MC, Zepka LQ, Jacob-Lopes E. Artificial lighting strategies in photobioreactors for bioenergy production by Scenedesmus obliquus CPCC05. SN Appl Sci [Internet]. 2019 [cited 2025 Aug 20];1:1695. 10.1007/s42452-019-1761-0

20. Herrera N, Echeverri F. Evidence of Quorum Sensing in Cyanobacteria by Homoserine Lactones: The Origin of Blooms. Water [Internet]. 2021 [cited 2023 Dec 19];13:1831. 10.3390/w13131831

21. Wu J, Keller DP, Oschlies A. Carbon dioxide removal via macroalgae open-ocean mariculture and sinking: an Earth system modeling study. Earth Syst Dynam [Internet]. 2023 [cited 2025 Sept 3];14:185–221. 10.5194/esd-14-185-2023

22. Yaakob MA, Mohamed RMSR, Al-Gheethi A, Aswathnarayana Gokare R, Ambati RR. Influence of Nitrogen and Phosphorus on Microalgal Growth, Biomass, Lipid, and Fatty Acid Production: An Overview. Cells [Internet]. 2021 [cited 2025 Aug 12];10:393. 10.3390/cells10020393

23. Chen Q, Chen Y, Xu Q, Jin H, Hu Q, Han D. Effective Two-Stage Heterotrophic Cultivation of the Unicellular Green Microalga Chromochloris zofingiensis Enabled Ultrahigh Biomass and Astaxanthin Production. Front Bioeng Biotechnol [Internet]. 2022 [cited 2023 Dec 19];10:834230. 10.3389/fbioe.2022.834230

24. Tripathi D, Singh M, Pandey-Rai S. Crosstalk of nanoparticles and phytohormones regulate plant growth and metabolism under abiotic and biotic stress. Plant Stress [Internet]. 2022 [cited 2023 Dec 19];6:100107. 10.1016/j.stress.2022.100107

25. Yang Z-Y, Huang K-X, Zhang Y-R, Yang L, Zhou J-L, Yang Q, et al. Efficient microalgal lipid production driven by salt stress and phytohormones synergistically. Bioresource Technology [Internet]. 2023 [cited 2023 Dec 19];367:128270. 10.1016/j.biortech.2022.128270

26. Lyu W, Zhang S, Xie Y, Chen R, Hu X, Wang B, et al. Effects of the exogenous N-acylhomoserine lactones on the performances of microalgal-bacterial granular consortia. Environmental Pollutants and Bioavailability [Internet]. 2022 [cited 2023 Dec 19];34:407–18. 10.1080/26395940.2022.2123046

27. Zhang Y, Zheng L, Wang S, Zhao Y, Xu X, Han B, et al. Quorum Sensing Bacteria in the Phycosphere of HAB Microalgae and Their Ecological Functions Related to Cross-Kingdom Interactions. IJERPH [Internet]. 2021 [cited 2023 Dec 19];19:163. 10.3390/ijerph19010163

28. Ei E, Park HH, Kuk YI. Soil Spraying of Water Extracts as a Plant Biostimulant Enhances Growth and Secondary Metabolites in Different Stages of Vegetable and Rice Crops [Internet]. 2024 [cited 2025 May 28]. 10.20944/preprints202411.1133.v1

29. Srinivasan P, Raja HD, Tamilvanan R. Effect of coconut water and cytokinins on rapid micropropagation of Ranunculus wallichianus Wight & Arnn—a rare and endemic medicinal plant of the Western Ghats, India. In Vitro CellDevBiol-Plant [Internet]. 2021 [cited 2025 May 28];57:365–71. 10.1007/s11627-020-10137-1

30. Jaubert M, Bouly J-P, Ribera d’Alcalà M, Falciatore A. Light sensing and responses in marine microalgae. Current Opinion in Plant Biology [Internet]. 2017 [cited 2025 Apr 13];37:70–7. 10.1016/j.pbi.2017.03.005

31. Chen Y, Xu C. How to narrow the CO2 gap from growth-optimal to flue gas levels by using microalgae for carbon capture and sustainable biomass production. Journal of Cleaner Production [Internet]. 2021 [cited 2025 Apr 13];280:124448. 10.1016/j.jclepro.2020.124448

32. Chutimanukul P, Sukdee S, Prajuabjinda O, Thepsilvisut O, Panthong S, Ehara H, et al. Exogenous Application of Coconut Water to Promote Growth and Increase the Yield, Bioactive Compounds, and Antioxidant Activity for Hericium erinaceus Cultivation. Horticulturae [Internet]. 2023 [cited 2025 Sept 3];9:1131. 10.3390/horticulturae9101131

33. Luligo-Montealegre WE, Prado-Alzate S, Ayala-Aponte A, Tirado DF, Serna-Cock L. Aloe vera Cuticle: A Promising Organic Water-Retaining Agent for Agricultural Use. Horticulturae [Internet]. 2024 [cited 2025 Sept 3];10:797. 10.3390/horticulturae10080797

34. Kumar S, Kalita S, Basumatary IB, Kumar S, Ray S, Mukherjee A. Recent advances in therapeutic and biological activities of Aloe vera. Biocatalysis and Agricultural Biotechnology [Internet]. 2024 [cited 2025 Jan 14];57:103084. 10.1016/j.bcab.2024.103084

35. Maira Batool, Malaika Khan, Maria Mubarak, Adil Hussain, Muafia Shafiq, Shamma Firdous, et al. A Wonder Plant Aloe vera L. (Liliaceae): An Overview of its Folk Traditional Uses, Phytoconstituents, Biological Activities, and Cosmaceutical Applications. PPASB [Internet]. 2023 [cited 2025 Jan 14];60. 10.53560/PPASB(60-3)857

36. Rawat A, Saxena A. Short review on composition and medicinal applications of aloe-vera. Mumbai, India; 2023 [cited 2025 Jan 14]. p. 030007. 10.1063/5.0141304

37. Rethinam P, Krishnakumar V. Composition, Properties and Reactions of Coconut Water. Coconut Water [Internet]. Cham: Springer International Publishing; 2022 [cited 2025 Jan 14]. p. 77–138. 10.1007/978-3-031-10713-9_4

38. Bastos CRV, Maia IB, Pereira H, Navalho J, Varela JCS. Optimisation of Biomass Production and Nutritional Value of Two Marine Diatoms (Bacillariophyceae), Skeletonema costatum and Chaetoceros calcitrans. Biology [Internet]. 2022 [cited 2025 Sept 10];11:594. 10.3390/biology11040594

39. Palikrousis TL, Manolis C, Kalamaras SD, Samaras P. Effect of Light Intensity on the Growth and Nutrient Uptake of the Microalga Chlorella sorokiniana Cultivated in Biogas Plant Digestate. Water [Internet]. 2024 [cited 2025 Sept 10];16:2782. 10.3390/w16192782

40. Sharmila D, Suresh A, Indhumathi J, Gowtham K. Impact of various color filtered LED lights on microalgae growth, pigments and lipid production. European Journal of Biotechnology and Bioscience. 2018;6:1–7. https://www.researchgate.net/publication/328635312

41. Tayebati H, Pajoum Shariati F, Soltani N, Sepasi Tehrani H. Effect of various light spectra on amino acids and pigment production of *Arthrospira platensis* using flat-plate photobioreactor. Preparative Biochemistry & Biotechnology [Internet]. 2024 [cited 2025 Jan 14];54:1028–39. 10.1080/10826068.2021.1941102

42. Niizawa I, Leonardi RJ, Irazoqui HA, Heinrich JM. Light wavelength distribution effects on the growth rate of Scenedesmus quadricauda. Biochemical Engineering Journal [Internet]. 2017 [cited 2025 Jan 14];126:126–34. 10.1016/j.bej.2016.09.006

43. Sánchez-Saavedra MDP, Sauceda-Carvajal D, Castro-Ochoa FY, Molina-Cárdenas CA. The Use of Light Spectra to Improve the Growth and Lipid Content of Chlorella vulgaris for Biofuels Production. Bioenerg Res [Internet]. 2020 [cited 2025 Jan 14];13:487–98. 10.1007/s12155-019-10070-1

44. Zhong Y, Jin P, Cheng JJ. A comprehensive comparable study of the physiological properties of four microalgal species under different light wavelength conditions. Planta [Internet]. 2018 [cited 2025 Apr 29];248:489–98. 10.1007/s00425-018-2899-5

45. Eloka-Eboka AC, Inambao FL. Effects of CO 2 sequestration on lipid and biomass productivity in microalgal biomass production. Applied Energy [Internet]. 2017 [cited 2025 Jan 14];195:1100–11. 10.1016/j.apenergy.2017.03.071

46. Wang B, Li Y, Wu N, Lan CQ. CO2 bio-mitigation using microalgae. Appl Microbiol Biotechnol [Internet]. 2008 [cited 2025 Apr 13];79:707–18. 10.1007/s00253-008-1518-y

47. Esteves AF, Soares OSGP, Vilar VJP, Pires JCM, Gonçalves AL. The Effect of Light Wavelength on CO2 Capture, Biomass Production and Nutrient Uptake by Green Microalgae: A Step Forward on Process Integration and Optimisation. Energies [Internet]. 2020 [cited 2025 Apr 13];13:333. 10.3390/en13020333

48. Kumar MS, Hwang J-H, Abou-Shanab RAI, Kabra AN, Ji M-K, Jeon B-H. Influence of CO2 and light spectra on the enhancement of microalgal growth and lipid content. Journal of Renewable and Sustainable Energy [Internet]. 2014 [cited 2025 Apr 13];6:063107. 10.1063/1.4901541

49. Jazzar S, Berrejeb N, Messaoud C, Marzouki MN, Smaali I. Growth Parameters, Photosynthetic Performance, and Biochemical Characterization of Newly Isolated Green Microalgae in Response to Culture Condition Variations. Appl Biochem Biotechnol [Internet]. 2016 [cited 2025 Apr 13];179:1290–308. 10.1007/s12010-016-2066-z

